# Simultaneous detection and quantification of adenine nucleotides in mammalian cultured cells by HPLC

**DOI:** 10.1101/2025.08.06.668918

**Authors:** Beatriz Kopel, Fernanda Manso Prado, Sofia Lígia Guimarães Ramos, Rafael Dias de Moura, Paolo Di Mascio, Nícolas Carlos Hoch, Marisa Helena Gennari de Medeiros, Nadja Cristhina de Souza Pinto

## Abstract

Adenine nucleotides, including ATP, ADP, ADP-ribose, AMP, NAD^+^ and NADH, play central roles in cellular homeostasis and involved in multiple metabolic and signaling pathways. Owing to their broad functional relevance in cell biology, the accurate quantification of these metabolites is essential for diverse research areas such as bioenergetics, cell signaling and cancer biology. Several analytical methods have been described for the measurement of adenine nucleotides, ranging from enzymatic assays to mass spectrometry-based approaches. In this study, we developed a reverse-phase high-performance liquid chromatography (RP-HPLC) method with UV-Vis detection that enables the simultaneous quantification of ATP, ADP, ADP-ribose, AMP, NAD^+^ and NADH. This method is simple, sensitive within the physiological concentration range of all analytes, and capable of detecting biologically relevant changes in ATP, AMP and NAD^+^ levels induced by pharmacological treatments. Therefore, it presents an accessible and reliable alternative for quantification of these nucleotides in biological samples.

## Introduction

Adenine nucleotides play essential roles in biological systems, ranging from energy metabolism to cellular signaling. Adenosine triphosphate (ATP), through coupled reactions, is the primary driver of thermodynamically unfavorable processes in living systems and is therefore widely regarded as the “energy currency” of the cell^1,2^. In addition to its bioenergetic function, ATP serves as the main phosphate donor in kinase-catalyzed phosphorylation reactions that regulate numerous signaling pathways. Both ATP and adenosine diphosphate (ADP) can also function as signaling molecules, notably in purinergic signaling and related pathways^3^. Adenosine monophosphate (AMP), generated from ADP hydrolysis, accumulates under conditions of high energy demand and act as a central regulator of cellular energy sensing through activation of the AMP-activated protein kinase (AMPK) pathway^4^. AMPK, in turn, modulates a wide range of downstream processes, including glucose and lipid metabolism, cell growth, autophagy, and other cellular pathways^5^.

Oxidized nicotinamide adenine dinucleotide (NAD^+^) plays a pivotal role in cellular redox reactions, acting as a coenzyme in key metabolic pathways such as glycolysis, fatty acid oxidation, the citric acid cycle^6,7^. The intracellular ratio of NAD^+^ to its reduced form, NADH, is an important indicator of cellular redox state and a critical regulator of multiple cellular responses. NAD(P)H also represents a major sources of reducing equivalents for antioxidant systems, including the glutathione peroxidase cycle, which relies on NADP^+^, generated by phosphorylation of NAD^+ 8^, to regenerate enzyme activity^9^.

Beyond their canonical bioenergetic functions, adenine nucleotides are also critical for genomic stability. In response to DNA damage, poly(ADP-ribose) polymerase 1 (PARP1) utilizes NAD^+^ as a substrate to catalyze the formation of poly(ADP-ribose) (PAR) chains, in which ADP-ribose (ADPr) moieties are covalently attached to target proteins, concomitant with the release of nicotinamide^10^. PARP1 activity is strongly induced by DNA damage, particularly single-strand breaks, leading to PARylation of histones surrounding the damage site and facilitating the recruitment of the DNA repair machinery^11,12^. Following lesion resolution, PAR chains are degraded by hydrolases such as poly(ADP-ribose) glycohydrolase (PARG), generating free ADPr monomers that can be recycled into NAD^+^ by the NAD salvage pathway^13,14^.

Given their widespread and fundamental roles, the quantification of ATP, ADP and AMP in biological samples has been pursued since the early 1950s, primarily using enzymatic and chromatographic approaches. Early methods relied on ion-exchange chromatography with borate complexes, requiring extensive sample processing to separate the nucleotides prior to analysis^15,16^. In 1954, the first luminescence-bases assay for ATP quantification was introduced, shortly after the discovery that ATP stimulates light emission in firefly lantern extracts^17^. This approach was later refined through the use of purified luciferin/luciferase systems and enzymatic conversion of ADP and AMP to ATP, forming the basis of most commercial luciferase-based ATP quantification kits available today^18^.

More recently, analytical techniques such as reverse phase high performance liquid chromatography (RP-HPLC) and liquid chromatography tandem mass spectrometry (LC-MS/MS) have been developed to enable the simultaneous quantification of ATP, ADP and AMP in biological samples^19^. Although LC-MS/MS offers superior sensitivity and lower limits of quantification (LOQ), its use can be limited by challenges in chromatographic separation using MS-compatible mobile phases and by degradation of ATP and ADP to AMP in the ionization source, which can compromise accurate quantification of the latter^20^, despite successfully implementation reported in the literature^21^.

The simultaneous determination of NAD^+^ and NADH also presents analytical challenges due to their distinct stability profiles across different pHs. NAD^+^ is more stable under acidic conditions, whereas NADH is stabilized at alkaline pH, likely owing to proton-catalyzed autoxidation of the reduced form^22^. Consequently, rapid metabolic quenching during sample preparation is critical for accurate measurements^23^. Nevertheless, NAD^+^ and NADH have been successfully quantified using colorimetric assays^24^, fluorescence spectrophotometry^25^, and by LC-MS/MS-based methods^22^.

In contrast, analytical methods for the quantification of free ADPr remain limited, as studies have focused on its polymerized form, PAR. In biological samples, PAR is predominantly measured using antibody-based approaches^26^. Measurements of free ADPr are comparatively scarce and have largely been restricted to *in vitro* assays of PARP1 activity using HPLC^27^ or to LC-MS/MS-based quantification alongside NAD^+^ following acidic extraction from yeast^28^ or mammalian cells^20^.

In the present study, we developed a method for the simultaneous detection of six metabolically relevant adenine nucleotides - ATP, ADP, AMP, ADPr, NAD^+^, and NADH (Figure 1) – in a single biological sample using RP-HPLC coupled to UV-Vis detection. This approach reduces the number of experimental replicates required, significantly shortens sample preparation and analysis time, and improves overall experimental sustainability and usability. Moreover, HPLC represents a more cost-effective and accessible alternative to LC-MS/MS. Importantly, the method developed here achieves limits of detection compatible with physiological concentration ranges of all six metabolites in cultured human cells. Validation experiments using pharmacological agents known to alter concentrations of ATP, AMP and NAD+ levels further demonstrated that this method reliably detects biologically meaningful variations in adenine nucleotide concentrations.

**Figure 1.**
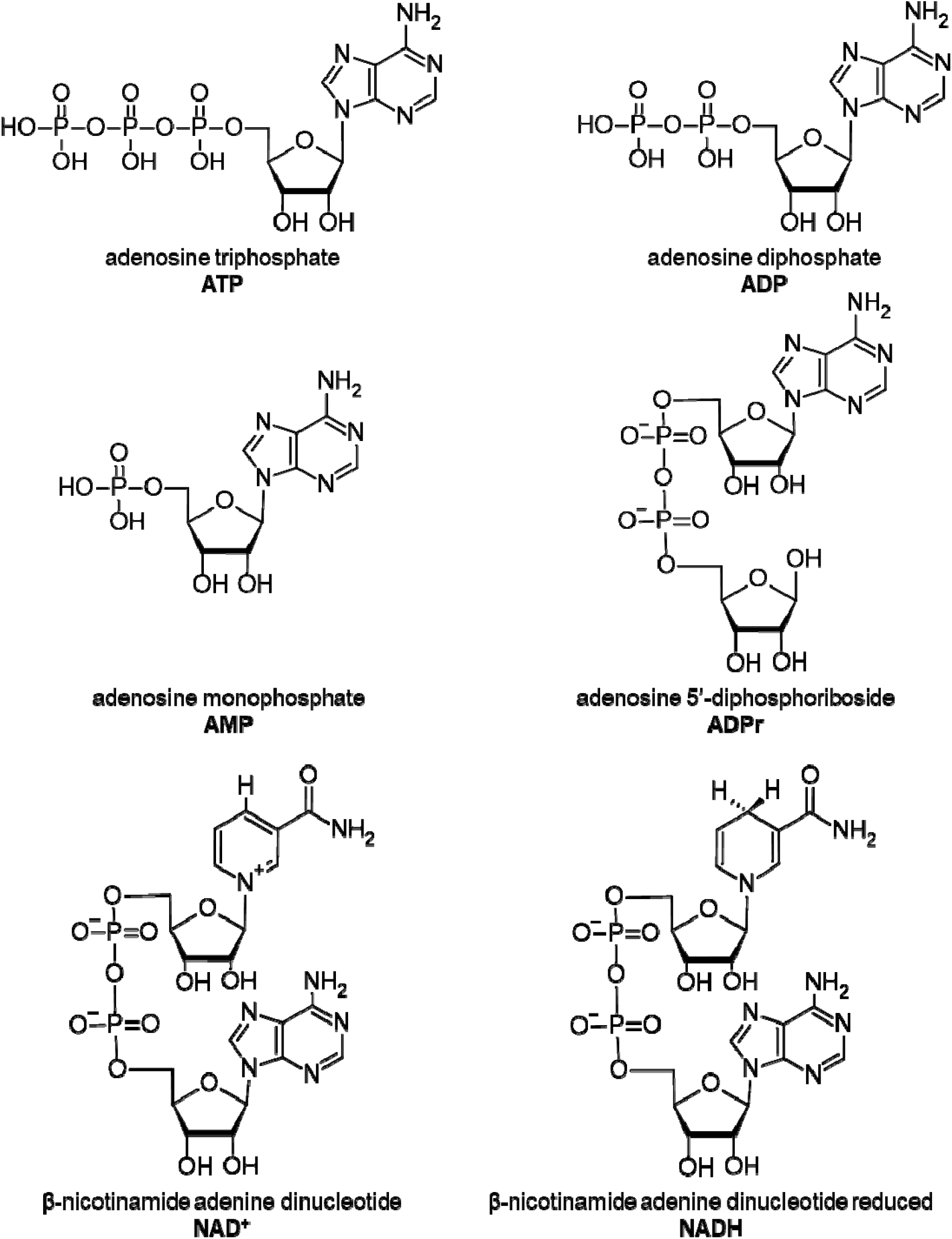
Chemical structure of the six adenine nucleotides analyzed here.

## Materials and methods

### 1. Cell culture and treatments

MRC-5 normal human pulmonary fibroblasts were maintained in DMEM high glucose containing 10% fetal bovine serum + 1% penicillin/streptomycin (complete medium) at 37 °C under 5% CO_2_ atmosphere. Cultures were routinely subcultured using 0.5% trypsin when 80% confluence was reached. For the treatments, cells were seeded in 100 mm plates and incubated with 20 µM carbonyl cyanide p-trifluoromethoxy phenylhydrazone (FCCP) for 60 min; 20 mM 2-deoxyglucose (2-deoxyGlu) + 1 µM oligomycin for 2.5 h; or 10 nM FK866 for 24 h. All treatments were carried out in complete medium, while control cells received 0.01% DMSO, as vehicle control.

### 2. Nucleotide extraction

Cells grown on 100 mm dishes were treated as described above and carefully washed with phosphate-buffer saline (PBS). After PBS removal, 1 mL of HPLC-grade methanol (Merck) containing 1 mM deferoxamine mesylate (Sigma-Aldrich) was added to the plate and homogenized with a cell scraper. Cell extracts were then transferred to a 2 mL microtube and incubated for two hours at 25 ºC with a 1,300-rpm agitation in a Thermomixer C (Eppendorf). Subsequently, samples were centrifuged at 20,000g for 10 minutes at 4 ºC. The supernatant, which contains all nucleotides, was stored at -80 ºC. The samples were dried at 20 ºC in a SpeedVac and then stored at – 80ºC until the analysis. The pellet was stored at – 20 ºC for protein quantification. Figure 2 depicts the experimental workflow for extraction and analysis of samples.

**Figure 2.**
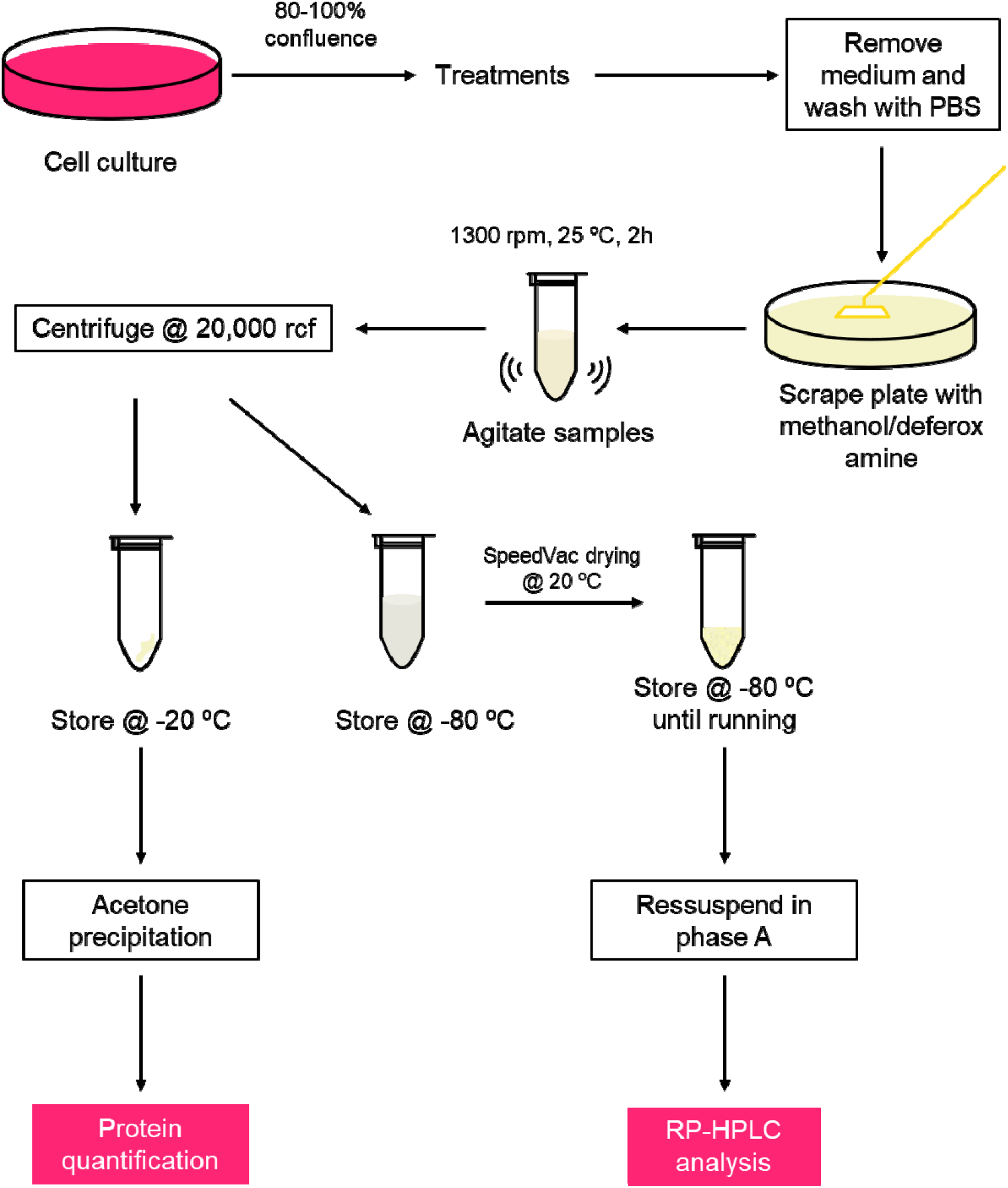
Schematic representation of the experimental workflow.

### 3. Protein purification and quantification

The pellets obtained after separation of the supernatant for nucleotide quantification were used for protein determination. The pellets were suspended in 250 µL of freshly prepared extraction buffer (100 mM ammonium bicarbonate, 8 M urea) and mixed in a Thermomixer at 25 ºC until completely dissolved. When pellet solubilization was not complete, samples were mixed at 1,000 rpm at 55 ºC in a Thermomixer for up to 1 hour. Subsequently, 1 mL of ice-cold acetone was added to the samples and vortexed thoroughly. The samples were kept at – 20 ºC overnight for protein precipitation followed by centrifugation at 13,000g for 10 minutes at 4 ºC. The supernatant was completely removed, and samples were air-dried in a fume hood. The dry pellets were resuspended in 200 µL of extraction buffer and used for protein quantification. Before quantification, samples were diluted so the final concentration of urea did not exceed 3 M, to avoid interfering in the protein quantification methods. We used Bio-Rad Protein Assay Dye Reagent Concentrate (Bio-Rad) according to the manufacturer’s instruction, and bovine gamma globulin (Bio-Rad) as standard.

### 4. RP-HPLC analysis

All analytes were analyzed in a single run using a Luna C18(2) 5 µm 100 Å 250 x 4.6 mm column (Phenomenex) with guard column AQ C18 4X3 mm ID (Phenomenex) in a SCL-40 HPLC System (Shimadzu). Two mobile phases were utilized: phase A being 25 mM diammonium hydrogen phosphate ((NH_4_)_2_HPO_4_) at pH 6,0, adjusted using 85% phosphoric acid, and phase B being HPLC-grade methanol. Column oven temperature was 25ºC and the flow was 0,8 mL/min. Samples were separated using the following gradient: 0-10 min 1% B, 10-15 min 1%-20% B, 15-20 min 20% B, 20-22 min 20%-40% B, 22-27 min 40% B, 27-30 min 40%-1% B, 30-40 min 1% B. Analytes were monitored at 254 nm. It should be noted that the analytes of interest elute before the 20-min running time, but the 40-min run ensures that the column is properly cleaned before the next sample is injected (Supplementary Figure S1).

Dried samples prepared as above were resuspended in 100 µL of phase A, agitated at 1,000 rpm for 10 min at 37 ºC and centrifuged at room temperature for 10 min at 10,000g. The supernatant was transferred to a new tube and 45 µL were injected per analysis. The same injection volume was also used to prepare the standard curves, which consisted of solutions in the 0.1 to 200 µg/mL concentration range for each analyte. The concentrations used for each analyte, in μM, are presented in Supplementary Table 7. We note that to ensure the reproducibility of the standard curves, all standards must be freshly prepared immediately before the experiment, as we observed that ATP and NADH, shown in Supplementary Figure S2, are rapidly degraded when stored at -20 ºC. All reagents used here were acquired from Sigma-Aldrich.

### 5. Recovery assay for estimation of analyte loss during the experimental procedure and limit of quantification estimations

To test how the extraction procedure affected analyte recovery and estimate the loss of each metabolite during the nucleotide extraction process, samples containing all analytical standards were submitted to similar processing as the biological samples and the peak areas obtained in RP-HPLC analysis were then compared to the area of control samples, which were not submitted to extraction. For this test, ATP, ADP, ADP-ribose and AMP were tested together to account for the possibility of nucleotide hydrolysis into AMP during the extraction process. NAD^+^ and NADH were evaluated separately. Loss rates were calculated as the percentage of peak area lost after extraction and used as correction factors in the calculation of the concentration for biological samples. The limit of quantification (LOQ) of each analyte was determined by visual inspection of the integrable peaks, with the lowest concentration with an integrable peak being considered the LOQ. The LOQ was estimated based on at least 3 independent experiments.

### 6. Statistical analysis

Linear regressions were obtained using GraphPad Prism 10 and Origin 2024. To evaluate statistical significance, one-way ANOVA with a Tukey post-hoc test for each metabolite was performed using GraphPad Prism 10.

## Results and discussion

### 1. Sample preparation procedure

Several methodologies have been reported for the preparation of samples for adenine nucleotide quantification, employing distinct strategies^19–21,28,29^. A fundamental aspect shared among these approaches is metabolic quenching, which is essential to preserve nucleotide concentrations as close as possible to their intracellular levels. Acidification is a commonly used quenching strategy, as it rapidly inactivates ATPases and thereby preserves ATP levels^30,31^. However, this approach is incompatible with experimental designs that require the simultaneous determination of NAD^+^ and NADH, because NADH is spontaneously oxidized to NAD^+^ under acidic conditions^22^, leading to distortion of their relative concentration.

To overcome this limitation, metabolic quenching can alternatively be achieved using organic solvents, such as methanol or combinations of solvents^32,33^, which was the strategy adopted in the present study. To preserve the cellular NAD^+^/NADH ratio during extraction, 100 mM deferoxamine mesylate, an iron chelating agent, was added to 100% methanol. This combination was found to effectively minimize metabolite loss during sample preparation (data not shown).

### 2. RP-HPLC provides a complete separation of six metabolites and low LOQ

Many biologically relevant processes lead to changes in absolute concentrations and/or relative ratios of adenine nucleotides, underscoring the importance of accurately quantifying these species. Ideally, all analytes should be measured in the same samples to minimize artifacts arising from sample handling and processing. Accordingly, we developed an HPLC-based protocol enabling the simultaneous determination of six adenine nucleotides with central roles in mammalian energy metabolism: ATP, ADP, AMP, NAD+, NADH and ADP-ribose. As shown in Figure 3, the chromatography conditions established herein provided adequate resolution of all six analytes.

**Figure 3.**
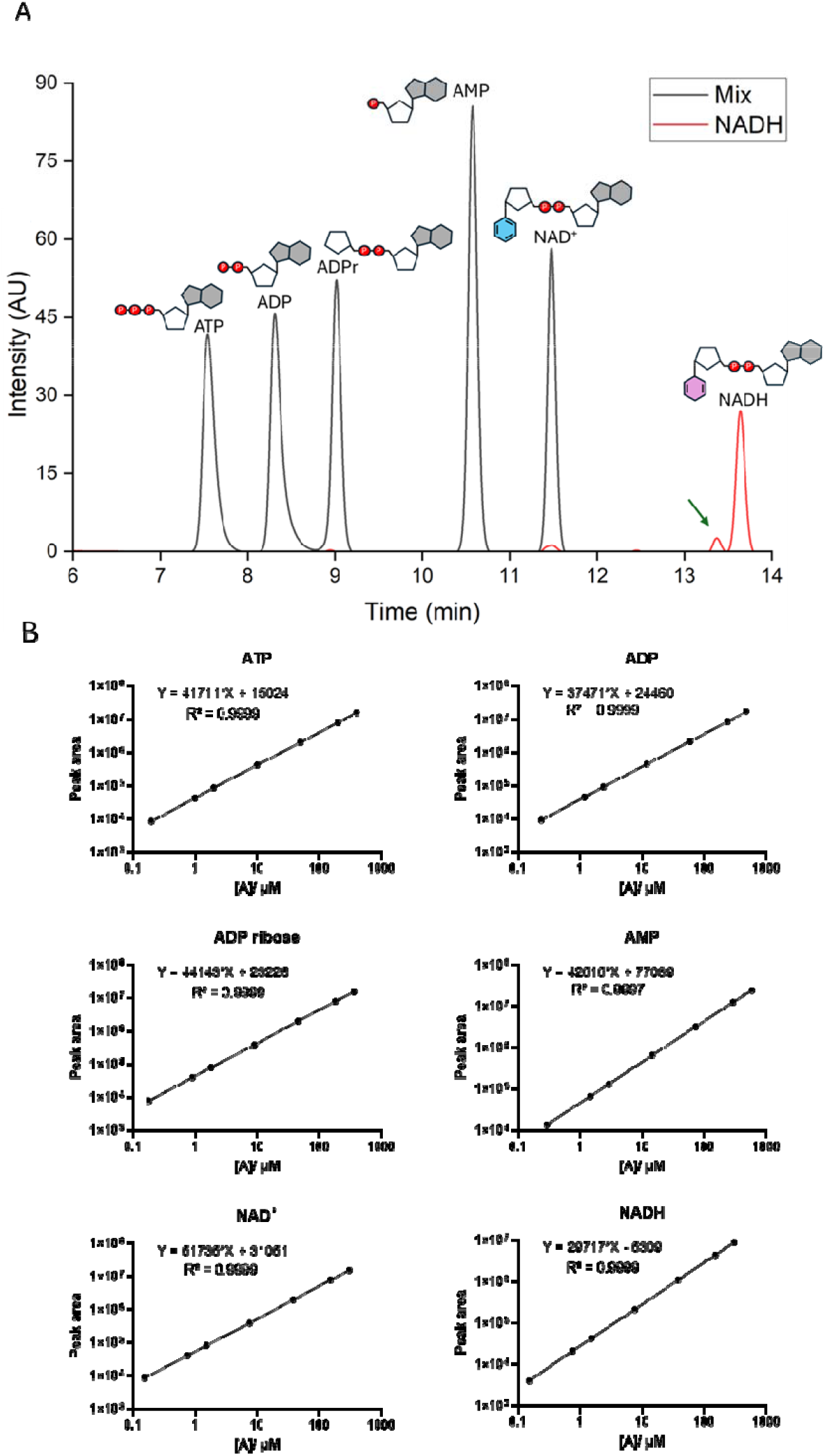
Separation profile of standards and linearity of standard curves. A: Chromatogram of standard analytes at 5 µg/mL concentration. ATP, ADP, ADPr, AMP and NAD+ were run together, presented as “Mix” curve, and NADH was run separately (red). NAD+ formed due to NADH degradation is indicated by the green arrow. B: Standard curves for each analyte obtained in the conditions shown in A. Equations are represented as y = ax + b and both axis for linear regressions are represented on a logarithmic scale.

It should be noted that the standard chromatograms shown in Figure 3A were obtained using solutions containing all nucleotides except NADH. NADH standards were analyzed separately to prevent interferences with NAD^+^ quantification caused by artifactual oxidation of NADH present in mixed standards. Even under these conditions, a small peak corresponding to the NAD^+^ retention time was detected in the NADH standard chromatogram (Figure 3, green arrow, red line), illustrating the susceptibility of NADH to oxidation. Figure 3 shows the individual standard curves for all six nucleotides.

Using these standards, we determined both the analyte-specific loss rates associated with experimental handling and the limits of quantification (LOQ) for each compound. ATP, ADP, ADP-ribose and NAD^+^ exhibited comparable loss rates, with a mean loss of 16.82% loss (Table 1). This parameter is particularly relevant for biological samples, as extensive analyte loss during sample preparation may preclude quantification if post-extraction concentrations fall below the LOQ. Taken together, the relatively low loss rate and LOQ for these nucleotides indicate that this method is both sensitive and well suited for biological applications.

**Table 1:** Retention time, limit of quantification (LOQ), and estimation of loss due to the sample preparation procedure for each analyte.

|  | ATP | ADP | ADPr | AMP | NAD<sup>+</sup> | NADH |
| --- | --- | --- | --- | --- | --- | --- |
| Retention time (min) | 7,5 | 8,4 | 9,1 | 10,7 | 11,5 | 13,7 |
| LOQ (nM) | 197 | 234 | 179 | 288 | 151 | 150 |
| Loss rate (%) | 15.25 | 16.45 | 20.25 | -18.70 | 15.29 | 30.27 |

The LOQ is a critical performance metric for analytical methods applied to biological samples, given that endogenous metabolite concentrations may approach the detection limits. As summarized in Table 1, the LOQ values obtained with the optimized HPLC method ranged from 150 to 288 nM, which is compatible with the analysis of biological samples, in which most of the metabolites are typically present in millimolar range^34^.

In contrast, AMP and NADH displayed loss rates that differed from those of the other nucleotides (Table 1). In mixtures containing ATP, ADP, ADP-ribose, and AMP, higher than expected levels of AMP were detected after extraction, resulting in a negative apparent loss rate. This increase is likely attributable to spontaneous hydrolysis of ATP and ADP to AMP during sample processing. NADH exhibited a higher loss rate (30.27%) than the other metabolites, which can be partially explained by its oxidation to NAD^+^. Collectively, these findings highlight the importance of determining loss rates for each individual analyte and experimental protocol and emphasize that loss should not be considered an intrinsic property of the extraction method alone, but rather as a variable dependent on the chemical properties and reactivity of the metabolites analyzed.

### 3. Validation experiments with cultured human cells detect expected biological responses

To validate the ability of the present method to detect biologically relevant changes in adenine nucleotide concentrations, MRC-5 cells were subjected to experimental conditions known to modulate levels of the analytes of interest. Cells were treated with the nicotinamide phosphoribosyl transferase (NAMPT) inhibitor FK866 ^35^, which impairs the NAD^+^ salvage pathway and thereby reduced intracellular NAD^+^ levels. A second condition involved treatment with the oxidative phosphorylation uncoupler FCCP, which stimulates mitochondrial respiration and is expected to decrease NADH levels. A third condition consisted of combined treatment with oligomycin, an inhibitor of the ATP synthase, and 2-deoxyglucose, a potent inhibitor of glycolysis; together, these agents are expected to decrease ATP levels and concomitantly increase AMP concentration. Samples were processed as described above, and representative chromatograms of the treated cells are shown in Figure 4A. Visual inspection of peak intensities revealed clear treatment-dependent changes in metabolite levels, which are quantitatively presented in Figures 4B-G.

**Figure 4.**
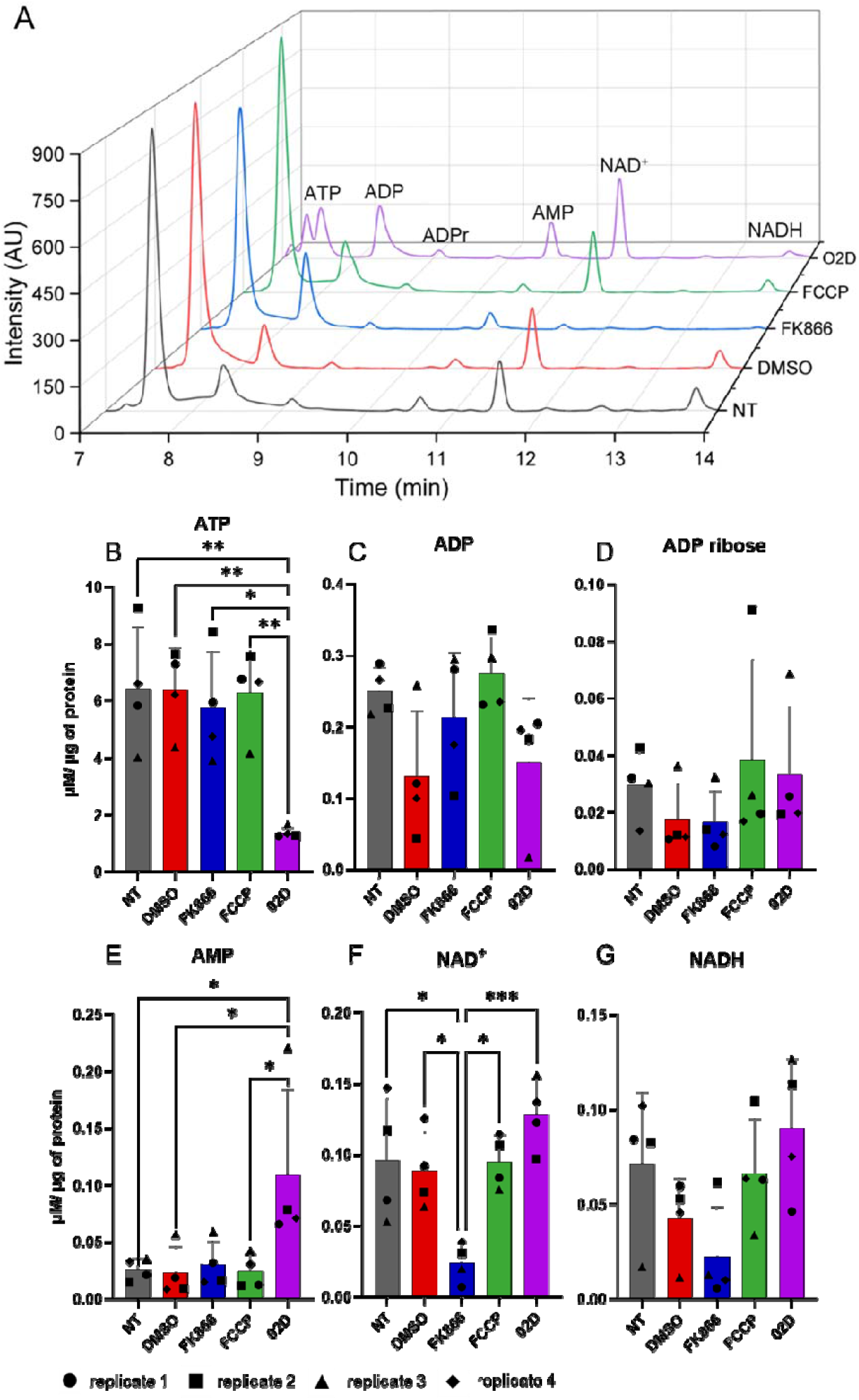
Simultaneous quantification of six adenine nucleotides in biological samples. A: representative chromatograms for each treatment, where NT: non-treated control; DMSO: 0.01% DMSO for 24h; FK866: 10 nM FK866 for 24h; FCCP: 20 µM FCCP for 2h; O2D: 1 µM oligomycin + 20 mM 2-deoxyglucose for 2h30min. B, C, D, E, F, G: concentration of each adenine nucleotide, normalized by protein mass in the extracts (µM/ µg of total protein). Results are presented as mean ± standard deviation of four independent biological replicates, performed in duplicate. Different biological replicates are represented as different geometrical forms, indicated at the bottom of the figure.

As anticipated, FK866 treatment resulted in a significant reduction in NAD^+^ concentration (Figure 4F; p < 0.05 relative to both control conditions), accompanied by a modest, non-significant decrease in NADH levels (Figure 4G). Despite these changes, the NAD^+^/NADH ratio was not significantly altered following FK866 treatment (Figure 4J), due to the concomitant decrease in both metabolites. These findings underscore the importance of quantifying the absolute concentration of individual nucleotides, rather than relying solely on metabolite ratios.

The results obtained following combined oligomycin and 2-deoxyglucose (O2D) treatment further support the notion that ratios may fail to capture biologically meaningful alterations. Although statistically significant changes in ATP (Figure 4B) and AMP (Figure 4E) concentrations were observed, neither the ATP/ADP nor the ATP/AMP ratios differed significantly between treated and control samples (Figures 5A and B). In the case of the ATP/AMP ratio (Figure 5B), relatively high sample variability likely contributed to the lack of statistical significance.

**Figure 5.**
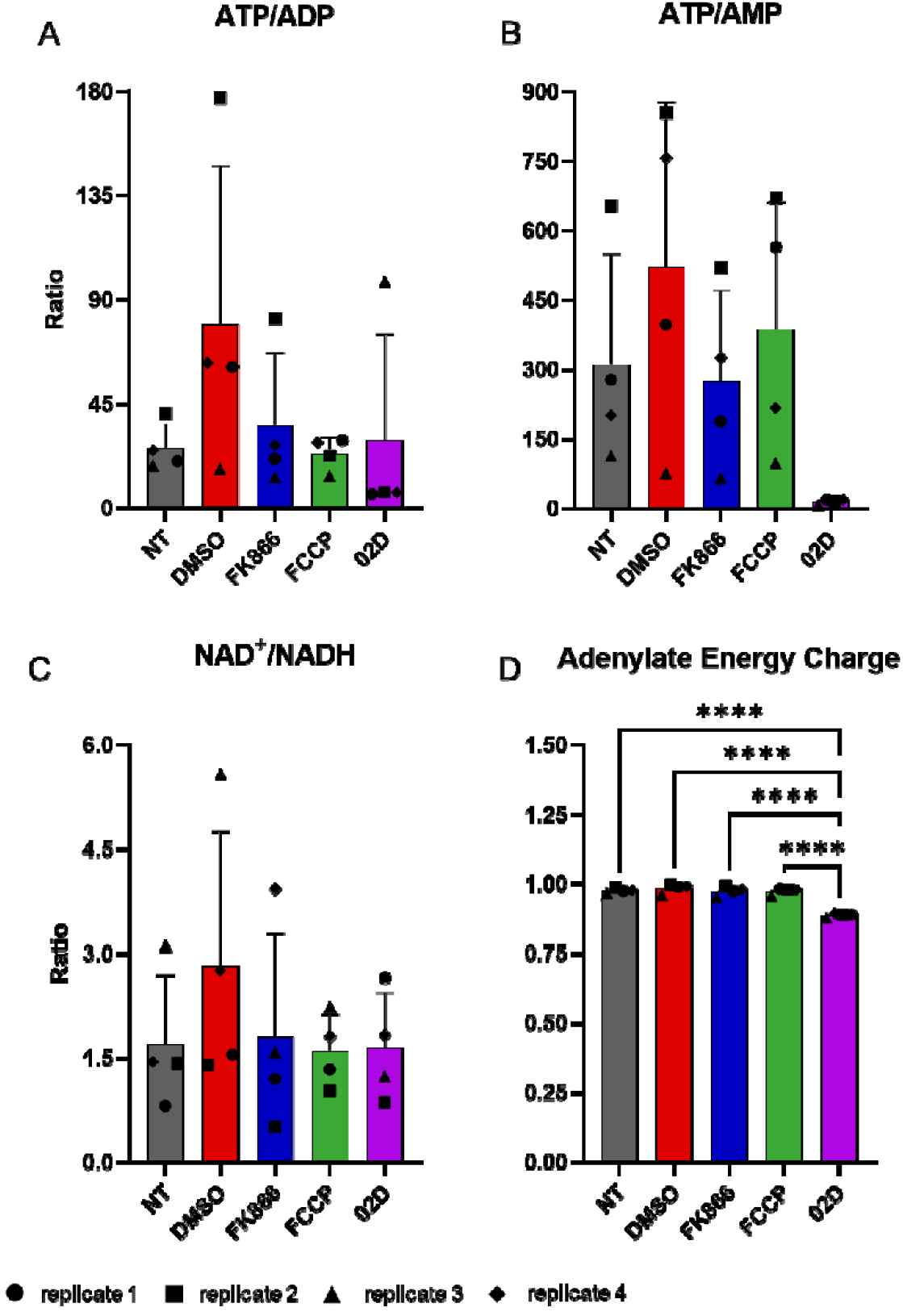
Ratios of adenine nucleotides calculated from the data in Figure 4 B-G. A-C: Usual ATP/ADP, ATP/AMP and NAD+/NADH ratios. D: Adenylate energy charge (AEC), calculated as (ATP + 0.5 ADP)/(ATP + ADP + AMP). ). Results are presented as mean ± standard deviation of four independent biological replicates, performed in duplicate. Different biological replicates are represented as different geometrical forms, indicated at the bottom of the figure.

The adenylate energy charge (AEC), a well-established indicator of cellular metabolic state and ATP-dependent processes, was also calculated for all conditions. AEC values range from 0 to 1^36^, and were determined using the equation (ATP + 0.5 ADP)/(ATP + ADP + AMP). Across the conditions analyzed, AEC remained largely stable (Figure 5D), with statistically significant differences observed between the O2D-treated sample and all other conditions.

## Conclusion

Given the central role of adenine nucleotides in biological processes, robust analytical methods are required for their accurate quantification in biological samples. Nonetheless, the applicability of several existing approaches is constrained by the high salt concentrations employed during sample preparation, as well as the need. For specialized instrumentation, such as mass spectrometry, which is not readily available to many cell biology laboratories. In this context, we developed an improved method based on HPLC with diode array detector (HPLC-DAD) that enables the simultaneous analysis of six adenine nucleotides in a single chromatographic run. The method relies on a simple and rapid sample preparation procedure that allows samples to be dried and stored, thereby decoupling biological experimentation from chromatographic analyses and permitting parallel processing, which enhances reproducibility. Furthermore, the use of organic solvent extraction in combination with deferoxamine minimizes artifactual degradation of analytes, thereby improving the accuracy of quantification. Simultaneous determination of all six adenine nucleotides within the same sample also contributes to reduced inter-sample variability and increased analytical precision.

We further demonstrated that this method is sufficiently sensitive to detect biologically relevant changes in nucleotide levels in cultured mammalian cells, with average intracellular concentrations well above the limit of detection for each analyte. Experiments employing established pharmacological interventions targeting cellular bioenergetics underscore the importance of measuring absolute nucleotide concentrations, as biologically meaningful effects may not be apparent when only metabolite ratios are considered. Notably, although FK866 treatment did not significantly alter the NAD^+^/NADH ratio, absolute quantification revealed a concomitant decrease in both NAD^+^ and NADH levels, an effect that would have remained undetected if only ratios had been assessed.

Taken together, the simplicity, adaptability and sensitivity of the sample preparation and chromatography procedures suggest that this method represents an accessible and reliable alternative to LC-MS/MS-based approaches for the quantitation of adenine nucleotides in biological samples of diverse origins.

## Acknowledgements

This work was supported by Fundação de Amparo à Pesquisa do Estado de São Paulo (FAPESP) grants 2024/07910-7 to NCS-P, CEPID Redoxoma 2013/07937–8 to PDM and MHGM, and 2018/18007-5 to NCH. BK is the recipient of FAPESP fellowship 2022/16432-6, RDM is the recipient of FAPESP fellowship 2020/02701-0, and SLRG is supported by Coordenação de Aperfeiçoamento de Pessoal de Nível Superior (CAPES). MHGM and PDM are supported by Universidade de São Paulo (USP) NAP Redoxoma (PRPUSP 2011.1.9352.1.8). NCS-P is supported by Conselho Nacional de Desenvolvimento Científico e Tecnológico (CNPq) grant 311588/2021-2, MHGM is supported by CNPq grant 304945/2021-8, and PDM is supported by CNPq grant 304350/2023-0 and John Simon Guggenheim Memorial Foundation. We thank Adriana M. P. Wendel for technical assistance and Bianca S. Vargas for the support.

## Author contributions

BK, conceptualization, investigation, writing-original draft; FMP, investigation; RDM, investigation; SLGR, investigation; PDM, funding acquisition, writing-review and editing; NCH, funding acquisition, writing-review and editing; MHGM, funding acquisition, writing-review and editing; NCS-P, conceptualization, funding acquisition, supervision, writing-original draft.

## Supporting information

In the supporting information section, we included Tables S1-6 with the concentrations of ATP, ADP, AMP, ADP ribose, NAD^+^ and NADH in µM prior to normalization for each experimental condition in each biological replicate. We also included Table S7, that details the analyte concentrations in µM used to prepare the standard curves. Figure S1 is a representative chromatogram without temporal cut and Figure S2 illustrates the degradation of NADH standard after storage at -20ºC.

## For Table of Contents Only

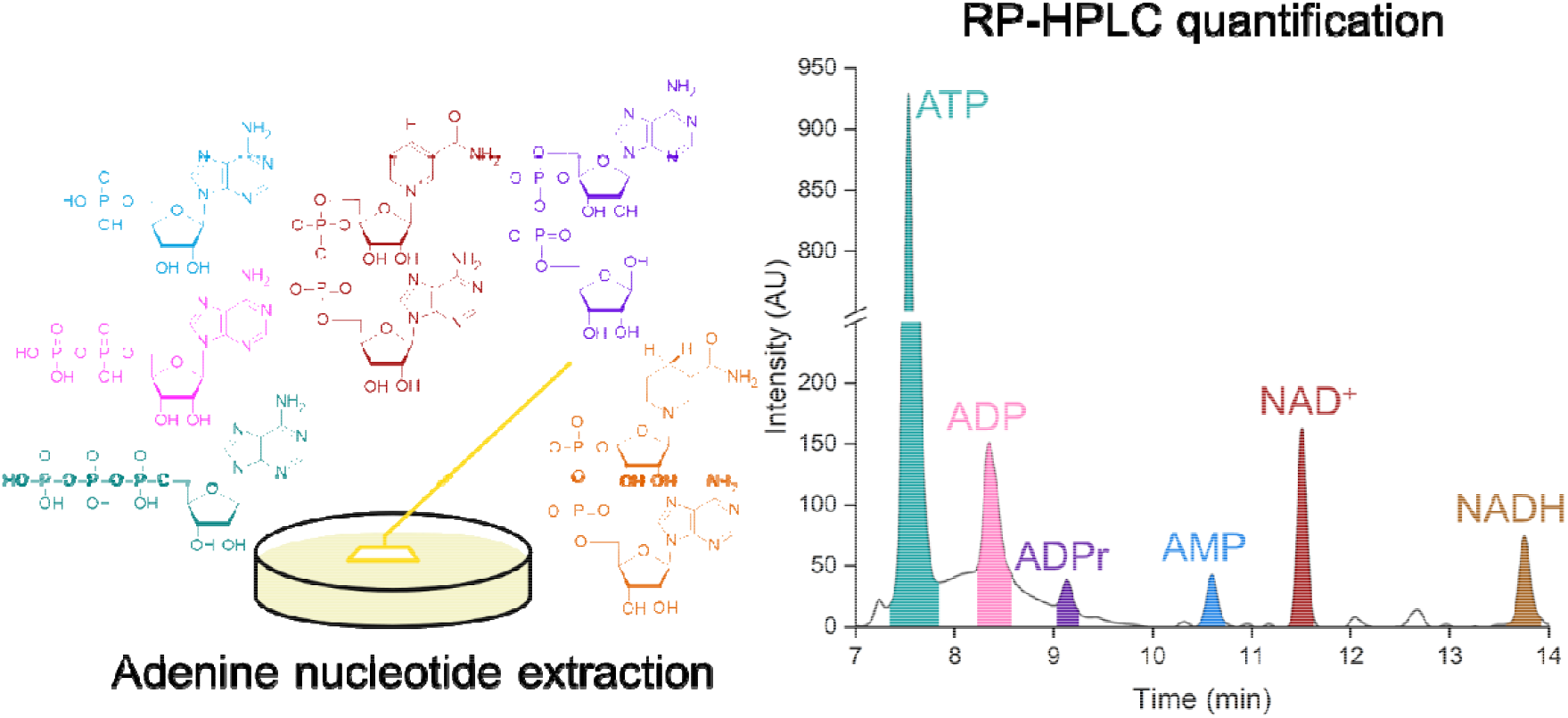

## References

(1) Nelson, D. L.; Cox, M. M.; Hoskins, A. A.; Lehninger, A. L. Lehninger Principles of Biochemistry, Eighth edition.; Macmillan International Higher Education: New York, 2021.

(2) Knowles, J. R. Enzyme-Catalyzed Phosphoryl Transfer Reactions. Annu. Rev. Biochem. 1980, 49 (1), 877–919. 10.1146/annurev.bi.49.070180.004305.

(3) Huang, Z.; Xie, N.; Illes, P.; Di Virgilio, F.; Ulrich, H.; Semyanov, A.; Verkhratsky, A.; Sperlagh, B.; Yu, S.-G.; Huang, C.; Tang, Y. From Purines to Purinergic Signalling: Molecular Functions and Human Diseases. Sig Transduct Target Ther 2021, 6 (1), 162. 10.1038/s41392-021-00553-z.

(4) Hardie, D. G. AMP-Activated Protein Kinase—an Energy Sensor That Regulates All Aspects of Cell Function. Genes Dev. 2011, 25 (18), 1895–1908. 10.1101/gad.17420111.

(5) Mihaylova, M. M.; Shaw, R. J. The AMPK Signalling Pathway Coordinates Cell Growth, Autophagy and Metabolism. Nat Cell Biol 2011, 13 (9), 1016–1023. 10.1038/ncb2329.

(6) Amjad, S.; Nisar, S.; Bhat, A. A.; Shah, A. R.; Frenneaux, M. P.; Fakhro, K.; Haris, M.; Reddy, R.; Patay, Z.; Baur, J.; Bagga, P. Role of NAD+ in Regulating Cellular and Metabolic Signaling Pathways. Molecular Metabolism 2021, 49, 101195. 10.1016/j.molmet.2021.101195.

(7) Cantó, C.; Menzies, K. J.; Auwerx, J. NAD+ Metabolism and the Control of Energy Homeostasis: A Balancing Act between Mitochondria and the Nucleus. Cell Metabolism 2015, 22 (1), 31–53. 10.1016/j.cmet.2015.05.023.

(8) Ying, W. NAD+ and NADH in Cellular Functions and Cell Death. Front Biosci 2006, 11 (1), 3129. 10.2741/2038.

(9) Pei, J.; Pan, X.; Wei, G.; Hua, Y. Research Progress of Glutathione Peroxidase Family (GPX) in Redoxidation. Front. Pharmacol. 2023, 14, 1147414. 10.3389/fphar.2023.1147414.

(10) Lüscher, B.; Ahel, I.; Altmeyer, M.; Ashworth, A.; Bai, P.; Chang, P.; Cohen, M.; Corda, D.; Dantzer, F.; Daugherty, M. D.; Dawson, T. M.; Dawson, V. L.; Deindl, S.; Fehr, A. R.; Feijs, K. L. H.; Filippov, D. V.; Gagné, J.; Grimaldi, G.; Guettler, S.; Hoch, N. C.; Hottiger, M. O.; Korn, P.; Kraus, W. L.; Ladurner, A.; Lehtiö, L.; Leung, A. K. L.; Lord, C. J.; Mangerich, A.; Matic, I.; Matthews, J.; Moldovan, G.; Moss, J.; Natoli, G.; Nielsen, M. L.; Niepel, M.; Nolte, F.; Pascal, J.; Paschal, B. M.; Pawłowski, K.; Poirier, G. G.; Smith, S.; Timinszky, G.; Wang, Z.; Yélamos, J.; Yu, X.; Zaja, R.; Ziegler, M. ADP-ribosyltransferases, an Update on Function and Nomenclature. The FEBS Journal 2022, 289 (23), 7399–7410. 10.1111/febs.16142.

(11) Caldecott, K. W. DNA Single-Strand Break Repair and Human Genetic Disease. Trends in Cell Biology 2022, 32 (9), 733–745. 10.1016/j.tcb.2022.04.010.

(12) Ying, S.; Chen, Z.; Medhurst, A. L.; Neal, J. A.; Bao, Z.; Mortusewicz, O.; McGouran, J.; Song, X.; Shen, H.; Hamdy, F. C.; Kessler, B. M.; Meek, K.; Helleday, T. DNA-PKcs and PARP1 Bind to Unresected Stalled DNA Replication Forks Where They Recruit XRCC1 to Mediate Repair. Cancer Research 2016, 76 (5), 1078–1088. 10.1158/0008-5472.CAN-15-0608.

(13) Barkauskaite, E.; Jankevicius, G.; Ladurner, A. G.; Ahel, I.; Timinszky, G. The Recognition and Removal of Cellular Poly( ADP-ribose) Signals. The FEBS Journal 2013, 280 (15), 3491–3507. 10.1111/febs.12358.

(14) Moura, R. D. D.; Mattos, P. D. D.; Valente, P. F.; Hoch, N. C. Molecular Mechanisms of Cell Death by Parthanatos: More Questions than Answers. Genet. Mol. Biol. 2024, 47 (suppl 1), e20230357. 10.1590/1678-4685-gmb-2023-0357.

(15) Strehler, B.; Totter, J. Determination of ATP and Related Compounds. Methods of biochemical analysis 1954, 1, 341–355.

(16) Khym, J. X.; Cohn, W. E. The Separation of Sugar Phosphates by Ion Exchange with the Use of the Borate Complex1. J. Am. Chem. Soc. 1953, 75 (5), 1153–1156. 10.1021/ja01101a043.

(17) McElroy, W. D. The Energy Source for Bioluminescence in an Isolated System. Proc. Natl. Acad. Sci. U.S.A. 1947, 33 (11), 342–345. 10.1073/pnas.33.11.342.

(18) Kimmich, G. A.; Randles, J.; Brand, J. S. Assay of Picomole Amounts of ATP, ADP, and AMP Using the Luciferase Enzyme System. Analytical Biochemistry 1975, 69 (1), 187–206. 10.1016/0003-2697(75)90580-1.

(19) Menegollo, M.; Tessari, I.; Bubacco, L.; Szabadkai, G. Determination of ATP, ADP, and AMP Levels by Reversed-Phase High-Performance Liquid Chromatography in Cultured Cells. Methods Mol Biol 2019, 1925, 223–232. 10.1007/978-1-4939-9018-4_19.

(20) Trammell, S. A.; Brenner, C. TARGETED, LCMS-BASED METABOLOMICS FOR QUANTITATIVE MEASUREMENT OF NAD + METABOLITES. Computational and Structural Biotechnology Journal 2013, 4 (5), e201301012. 10.5936/csbj.201301012.

(21) Hiefner, J.; Rische, J.; Bunders, M. J.; Worthmann, A. A Liquid Chromatography-Tandem Mass Spectrometry Based Method for the Quantification of Adenosine Nucleotides and NAD Precursors and Products in Various Biological Samples. Front. Immunol. 2023, 14, 1250762. 10.3389/fimmu.2023.1250762.

(22) Braidy, N.; Villalva, M. D.; Grant, R. NADomics: Measuring NAD+ and Related Metabolites Using Liquid Chromatography Mass Spectrometry. Life 2021, 11 (6), 512. 10.3390/life11060512.

(23) Airhart, S. E.; Shireman, L. M.; Risler, L. J.; Anderson, G. D.; Nagana Gowda, G. A.; Raftery, D.; Tian, R.; Shen, D. D.; O’Brien, K. D. An Open-Label, Non-Randomized Study of the Pharmacokinetics of the Nutritional Supplement Nicotinamide Riboside (NR) and Its Effects on Blood NAD+ Levels in Healthy Volunteers. PLoS ONE 2017, 12 (12), e0186459. 10.1371/journal.pone.0186459.

(24) Bernofsky, C.; Swan, M. An Improved Cycling Assay for Nicotinamide Adenine Dinucleotide. Analytical Biochemistry 1973, 53 (2), 452–458. 10.1016/0003-2697(73)90094-8.

(25) Mayevsky, A.; Rogatsky, G. G. Mitochondrial Function in Vivo Evaluated by NADH Fluorescence: From Animal Models to Human Studies. Am J Physiol Cell Physiol 2007, 292 (2), C615–640. 10.1152/ajpcell.00249.2006.

(26) Kawamitsu, H.; Hoshino, H.; Okada, H.; Miwa, M.; Momoi, H.; Sugimura, T. Monoclonal Antibodies to Poly(Adenosine Diphosphate Ribose) Recognize Different Structures. Biochemistry 1984, 23 (16), 3771–3777. 10.1021/bi00311a032.

(27) Rudolph, J.; Roberts, G.; Muthurajan, U. M.; Luger, K. HPF1 and Nucleosomes Mediate a Dramatic Switch in Activity of PARP1 from Polymerase to Hydrolase. eLife 2021, 10, e65773. 10.7554/eLife.65773.

(28) Tong, L.; Lee, S.; Denu, J. M. Hydrolase Regulates NAD+ Metabolites and Modulates Cellular Redox. Journal of Biological Chemistry 2009, 284 (17), 11256–11266. 10.1074/jbc.M809790200.

(29) Menezes-Filho, S. L.; Amigo, I.; Prado, F. M.; Ferreira, N. C.; Koike, M. K.; Pinto, I. F. D.; Miyamoto, S.; Montero, E. F. S.; Medeiros, M. H. G.; Kowaltowski, J. Caloric Restriction Protects Livers from Ischemia/Reperfusion Damage by Preventing Ca2+-Induced Mitochondrial Permeability Transition. Free Radical Biology and Medicine 2017, 110, 219–227. 10.1016/j.freeradbiomed.2017.06.013.

(30) Vives-Bauza, C.; Yang, L.; Manfredi, G. Assay of Mitochondrial ATP Synthesis in Animal Cells and Tissues. In Methods in Cell Biology; Elsevier, 2007; Vol. 80, pp 155–171. 10.1016/S0091-679X(06)80007-5.

(31) Menezes-Filho, S. L.; Amigo, I.; Prado, F. M.; Ferreira, N. C.; Koike, M. K.; Pinto, I. F. D.; Miyamoto, S.; Montero, E. F. S.; Medeiros, M. H. G.; Kowaltowski, J. Caloric Restriction Protects Livers from Ischemia/Reperfusion Damage by Preventing Ca2+-Induced Mitochondrial Permeability Transition. Free Radical Biology and Medicine 2017, 110, 219–227. 10.1016/j.freeradbiomed.2017.06.013.

(32) Fritsche-Guenther, R.; Bauer, A.; Gloaguen, Y.; Lorenz, M.; Kirwan, J. A. Modified Protocol of Harvesting, Extraction, and Normalization Approaches for Gas Chromatography Mass Spectrometry-Based Metabolomics Analysis of Adherent Cells Grown Under High Fetal Calf Serum Conditions. Metabolites 2019, 10 (1), 2. 10.3390/metabo10010002.

(33) Abdel Rahman, A. M.; Pawling, J.; Ryczko, M.; Caudy, A. A.; Dennis, J. W. Targeted Metabolomics in Cultured Cells and Tissues by Mass Spectrometry: Method Development and Validation. Analytica Chimica Acta 2014, 845, 53–61. 10.1016/j.aca.2014.06.012.

(34) Milo, R.; Phillips, R. Cell Biology by the Numbers; Garland Science, Taylor & Francis Group: New York, NY, 2016.

(35) Hasmann, M.; Schemainda, I. FK866, a Highly Specific Noncompetitive Inhibitor of Nicotinamide Phosphoribosyltransferase, Represents a Novel Mechanism for Induction of Tumor Cell Apoptosis. Cancer Research 2003, 63 (21), 7436–7442.

(36) Henderson, J. F.; Paterson, A. R. P. FUNCTIONS OF NUCLEOTIDES. In Nucleotide Metabolism; Elsevier, 1973; pp 28–56. 10.1016/B978-0-12-340550-0.50008-3.

