## supplementary tables and figures for "Simultaneous detection and quantification of adenine nucleotides in mammalian cultured cells by HPLC"

Table S1: ATP concentrations in µM prior to normalization.

| Replicate | NT | DMSO | FK866 | FCCP | 02D |
| --- | --- | --- | --- | --- | --- |
| N1 | 2049,3 | 2568,9 | 2145,6 | 2845,4 | 431,4 |
| N2 | 4531,7 | 4772,2 | 4653,5 | 4112,2 | 601,3 |
| N3 | 1148,1 | 1049,3 | 806,9 | 1077,2 | 117,5 |
| N4 | 2313,6 | 2361,1 | 1823,2 | 2107,3 | 400,5 |

Table S2: ADP concentrations in µM prior to normalization.

| Replicate | NT | DMSO | FK866 | FCCP | 02D |
| --- | --- | --- | --- | --- | --- |
| N1 | 101,4 | 42,1 | 100,2 | 94,7 | 71,9 |
| N2 | 106,8 | 26,9 | 56,8 | 182,3 | 87,6 |
| N3 | 61,4 | 62,6 | 62,6 | 65,3 | 24,8 |
| N4 | 92,3 | 37,5 | 67,1 | 74,1 | 59,9 |

Table S3: ADP ribose concentrations in µM prior to normalization.

| Replicate | NT | DMSO | FK866 | FCCP | 02D |
| --- | --- | --- | --- | --- | --- |
| N1 | 11,3 | 3,7 | 2,9 | 7,9 | 9,1 |
| N2 | 19,8 | 7,4 | 7,7 | 49,5 | 9,3 |
| N3 | 8,5 | 8,9 | 7,1 | 6,7 | 5,2 |
| N4 | 4,8 | 4,3 | 4,6 | 5,3 | 6,0 |

Table S4: AMP concentrations in µM prior to normalization.

| Replicate | NT | DMSO | FK866 | FCCP | 02D |
| --- | --- | --- | --- | --- | --- |
| N1 | 7,4 | 6,5 | 11,3 | 5,0 | 23,0 |
| N2 | 6,9 | 5,6 | 9,0 | 6,1 | 37,6 |
| N3 | 9,9 | 13,8 | 12,7 | 11,0 | 17,2 |
| N4 | 11,5 | 3,1 | 5,6 | 9,7 | 21,7 |

Table S5: NAD^+^ concentration in µM prior to normalization.

| Replicate | NT | DMSO | FK866 | FCCP | 02D |
| --- | --- | --- | --- | --- | --- |
| N1 | 23,9 | 32,4 | 2,4 | 34,8 | 43,0 |
| N2 | 57,4 | 46,0 | 17,1 | 57,9 | 46,5 |
| N3 | 15,1 | 15,5 | 4,2 | 19,6 | 9,7 |
| N4 | 51,9 | 47,8 | 15,0 | 36,2 | 42,1 |

Table S6: NADH concentration in µM prior to normalization.

| Replicate | NT | DMSO | FK866 | FCCP | 02D |
| --- | --- | --- | --- | --- | --- |
| N1 | 29,6 | 21,0 | 2,0 | 26,6 | 15,8 |
| N2 | 47,8 | 32,9 | 34,0 | 56,7 | 54,4 |
| N3 | 4,9 | 2,7 | 2,4 | 8,8 | 9,7 |
| N4 | 35,9 | 17,4 | 3,8 | 20,0 | 23,3 |

Table S7: Analyte concentrations used in the standard curve, in μM

|  | ATP | ADP | ADPr | AMP | NAD+ | NADH |
| --- | --- | --- | --- | --- | --- | --- |
| C1 | 0,20 | 0,23 | 0,18 | 0,29 | 0,15 | 0,15 |
| C2 | 0,99 | 1,17 | 0,89 | 1,44 | 0,75 | 0,75 |
| C3 | 1,97 | 2,34 | 1,79 | 2,88 | 1,51 | 1,50 |
| C4 | 9,86 | 11,71 | 8,94 | 14,41 | 7,54 | 7,52 |
| C5 | 49,31 | 58,55 | 44,70 | 72,03 | 37,68 | 37,59 |
| C6 | 197,24 | 234,18 | 178,79 | 288,13 | 150,73 | 150,35 |
| C7 | 394,49 | 468,36 | 357,58 | 576,27 | 301,46 | 300,70 |

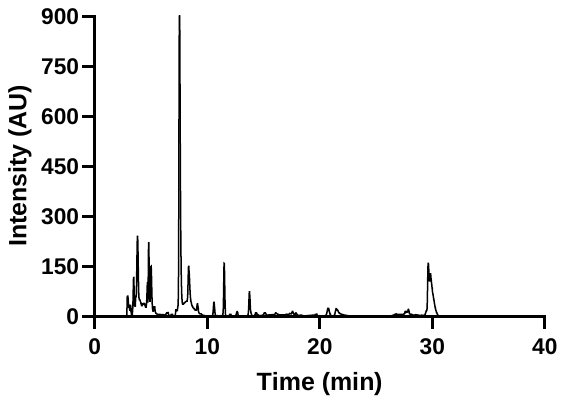

Figure S1: Representative chromatogram for the separation of an extract obtained from non-treated cells, under the same conditions described for the treated samples. The figure presents the entire chromatogram for the 40-minute running time.

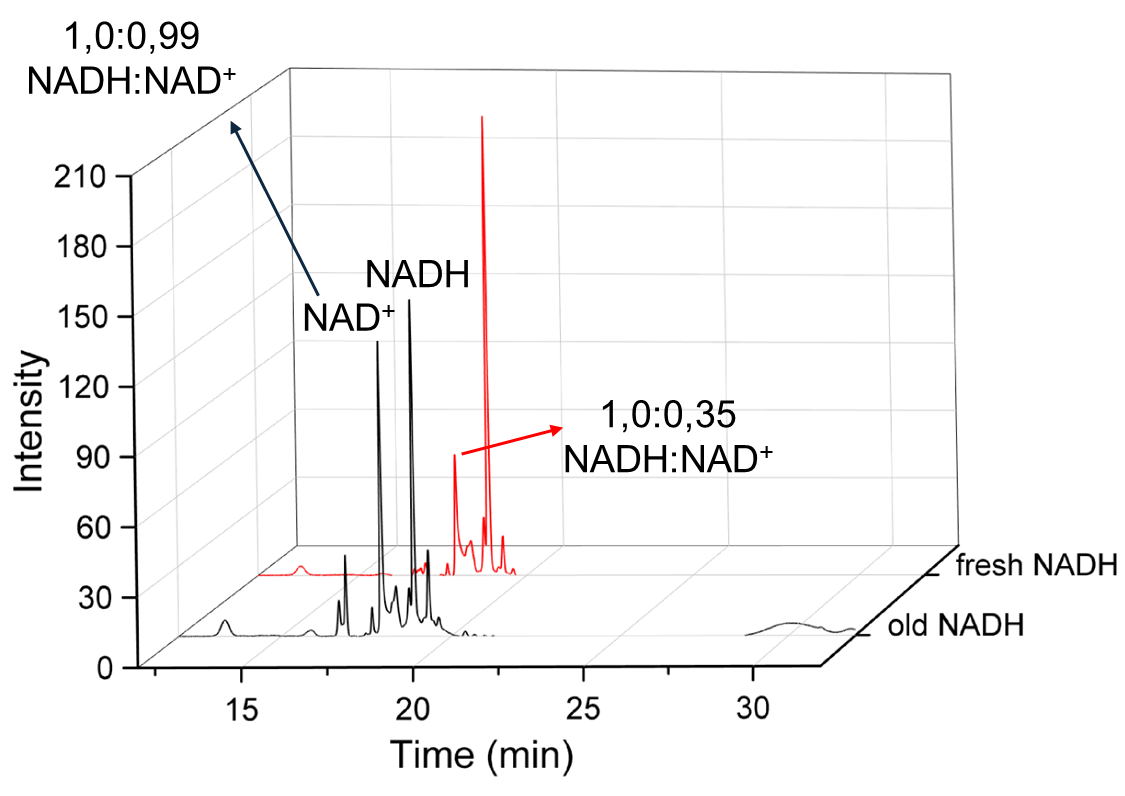

Figure S2: Degradation of NADH solutions stored at -20 ºC. Representative chromatograms of freshly made NADH standard solution (red line) and NADH standard solution of the same concentration (black line), stored for two months at -20 ºC. Both standard solutions were originally made at 50 µg/mL. Both arrows indicate the percentage of the NADH area that NAD+ peaks represent both in intact and degraded NADH standard.
